# Dynamics of CRISPR-mediated virus-host interactions in the human gut microbiome

**DOI:** 10.1101/2024.01.23.576851

**Authors:** Adrián López-Beltrán, João Botelho, Jaime Iranzo

**Author notes:** **Competing interests:** The authors declare no competing financial interests.

## Abstract

Arms races between mobile genetic elements and prokaryotic hosts are major drivers of ecological and evolutionary change in microbial communities. Prokaryotic defense systems such as CRISPR-Cas have the potential to regulate microbiome composition by modifying the interactions among bacteria, plasmids, and phages. Here, we used longitudinal metagenomic data from 130 healthy and diseased individuals to study how the interplay of genetic parasites and CRISPR-Cas immunity reflects on the dynamics and composition of the human gut microbiome. Based on the coordinated study of 80,000 CRISPR-Cas loci and their targets, we show that CRISPR-Cas immunity effectively modulates bacteriophage abundances in the gut. Acquisition of CRISPR-Cas immunity typically leads to a decrease in the abundance of lytic phages, but does not necessarily cause their complete disappearance. Much smaller effects are observed for lysogenic phages and plasmids. Conversely, phage-CRISPR interactions shape bacterial microdiversity by producing weak selective sweeps that benefit immune host lineages. Interestingly, distal (and chronologically older) regions of CRISPR arrays are enriched in spacers that are potentially functional and target crass-like phages and local prophages. This suggests that exposure to reactivated prophages and other endemic viruses is a major selective pressure in the gut microbiome that drives the maintenance of long-lasting immune memory.

## Introduction

Virus-host interactions are major determinants of community dynamics, composition, and evolution in microbial ecosystems (1–5). In two classic (and somewhat antagonistic) examples, phage outbreaks can produce the collapse of aquatic bacterial populations and cholera epidemics (6, 7), whereas sustained rates of phage-induced lysis positively affect ocean and soil communities by facilitating carbon and nitrogen cycling (8, 9). More recent studies have highlighted the potential of bacteriophages to shape host-associated microbiomes both directly, through phage-driven selective pressures, and indirectly, through the alteration of the host’s immune response (10–13).

Prokaryotes have developed a diverse repertoire of specialized defense mechanisms against viruses and other mobile genetic elements (MGE), such as plasmids and integrative and conjugative/mobilizable elements (ICE/IME), many of which have only recently started to be uncovered (14, 15). Among them, CRISPR-Cas systems are of particular interest as they provide their bearers with adaptive immunity (16–18) The adaptive nature of CRISPR-Cas systems is based on the acquisition of short DNA fragments from previously encountered MGE at the leader end of the CRISPR array. Such fragments, called spacers, serve as highly specific probes for the detection of the targeted MGE upon subsequent encounters. From a researcher’s perspective, CRISPR spacers constitute an ordered record that allow tracking virus-host interactions through time (19, 20).

Despite experimental studies in the laboratory have provided a general understanding of how CRISPR-Cas systems work, there are still many unknowns about their functioning in natural ecosystems (21). The most prominent question regards the actual effect that CRISPR-based immunity exerts on the composition and dynamics of microbial communities, both at the level of the virome and the bacterial or archaeal host population. Studies in free-living model ecosystems, such as hot springs, hypersaline lakes, and wastewater treatment plants, have demonstrated that CRISPR-Cas contributes to shaping a highly dynamic network of interactions among MGE and their prokaryotic hosts (19, 22–26). However, the paucity of appropriate datasets has hampered the application of similar methods to complex host-associated microbiomes. Some recurrent findings of previous studies are: (i) long-term coexistence of MGE with a polymorphic population of hosts that includes immune and sensitive lineages; (ii) high levels of local adaptation in the host CRISPR arrays; and (iii) high frequency of mutations in regions of MGE targeted by CRISPR spacers. Coexistence of MGE and host lineages is generally interpreted in terms of the diversity-promoting effects of clonal interference and herd immunity (27–31), whereas coevolution of CRISPR and MGE suggests that CRISPR-Cas is functional and the selective pressure that it imposes on MGE is strong enough to fuel mutational escape (32, 33). Interestingly, these conclusions may not extend to host-associated microbiomes. For example, single-sample measurements of *Prevotella* strains in the human gut and *Pseudomonas aeruginosa* in the lung display complete immunodominance of a single host lineage, possibly underscoring the occurrence of strong selective sweeps (34, 35).

A second major gap in our understanding of CRISPR-Cas systems concerns the time span of the immune memory, which results from the interplay of spacer gain and loss within the CRISPR array, mutation of the target MGE, and replacement of immune lineages in the population of hosts. Mathematical models predict that the size of CRISPR arrays is controlled by a trade-off between maintaining a large enough immune repertoire and avoiding the dilution effect of expressing too many spacers. Likewise, theoretical arguments suggest that the rates of spacer gain and loss must be finely tuned to keep CRISPR arrays sufficiently updated without losing immune memory too quickly (36, 37). These models operate on the basic assumption that distal spacers, generally assumed to be the oldest in the array (but see (38) for counterexamples) provide the host with long-term immune memory. However, the empirical and experimental evidence on this issue is inconclusive. On one hand, reduced transcription rates (39–41) and broad conservation of distal spacers across strains (42) suggest that those spacers do not play a relevant role in defense against local MGE. On the other hand, distal spacers often match conserved regions of viral genomes, which supports an active role of selection in maintaining these spacers for their higher potential in long-term immunity (43).

In this study, we leveraged a large longitudinal dataset from the integrative Human Microbiome Project (44) to investigate the joint dynamics of MGE and CRISPR-Cas immunity in the gut microbiome. We found evidence of effective (but incomplete) CRISPR-Cas immunity against lytic and lysogenic phages, short- and long-term coexistence of multiple MGE and host lineages, and selection for long-lasting immunity against endemic MGE provided by spacers from distal regions of CRISPR arrays.

## Methods

### Metagenomic data

Metagenomic and metatranscriptomic reads, assembled contigs, and taxonomic profiles from the Extended Human Microbiome Project – Inflammatory Bowel Disease (HMP2-IBD) were downloaded from https://ibdmdb.org. The metagenomic dataset includes 1,638 samples from 130 individuals, collected at 2-week intervals for approximately 1 year (44). Samples from participants with Crohn’s disease and ulcerative colitis were classified as dysbiotic or non-dysbiotic by applying the same criteria as in the original study.

### Assembly, annotation, and clustering of CRISPR arrays

For each sample of the HMP2-IBD dataset, we assembled putative CRISPR-Cas loci from metagenomic reads with the tool CasCollect v1.0 (45). CasCollect runs VSEARCH (46) in an iterative manner to identify reads that possibly belong to CRISPR-Cas loci and then assembles such reads with SPAdes (47). As baits for the first iteration of VSEARCH, we used the sequences of known CRISPR-associated genes (as provided by default by CasCollect) and an in-house database of CRISPR repeats that combined the default list of repeats provided by CasCollect, additional repeats from CRISPRdb (48) and Makarova et al. (49) clustered at 90% identity and coverage, and a set of previously unreported repeats obtained by running CRISPRCasTyper v1.8 (50) on the publicly available, preassembled contigs from the HMP2-IBD study. CasCollect was run with options --meta --noannotate --nucl. Putative CRISPR-Cas loci assembled by CasCollect were quality-filtered and annotated with CRISPRCasTyper v1.8, obtaining 79,475 high-confidence CRISPR arrays of which 4,282 (representing around 5% of the total arrays) included a Cas operon in their neighborhood. For comparison, we also searched for CRISPR-Cas loci using a “classical” sequential approach, consisting of whole-metagenome assembly followed by identification of CRISPR loci with CRISPRCasTyper. We tried this alternative approach with two assemblies: the one originally generated by the HMP2-IBD project (sample-level assembly with MEGAHIT, as described in (44)) and an in-house assembly with SPAdes at the individual level (coassembly of all the samples from the same individual). We found that the CRISPR arrays assembled with CasCollect were the most diverse in terms of repeats and spacers (Figure S1 and Table S1). Moreover, CasCollect produced more arrays of all possible lengths than the sample-level assembly approach (comparisons of array lengths with individual-level coassemblies are not informative because of possible chimaeras and implicit dereplication of arrays present in multiple samples). All in all, we concluded that CasCollect is more sensitive than whole-genome assembly-based methods and selected it for all downstream analyses.

Spacers and repeats from high-confidence CRISPR arrays were dereplicated using CD-Hit-Est v4.8.1 (51) with a 100% identity and coverage threshold and option *-g* 1 (slow and accurate mode), resulting in a total of 11,373 different repeats and 153,745 different spacers. The directionality of transcription in a subset of 4,054 arrays with adjacent Cas genes and >5 spacers was predicted with Cas Orientation (52), CRISPRLoci v1.0 (53), and CRISPRDirection v2.0 (54). Among these three prediction tools, Cas Orientation showed the best agreement with the expected drop in transcription for middle and distal spacers and was applicable to a larger fraction of arrays (Figure S2). Therefore, we chose Cas Orientation as the default method to present the results in the text and main figures. To explore the distribution of the number of spacers per array, we used the ’powerlaw’ python package (55).

A strict nesting criterion was applied to identify groups (or lineages) of highly similar CRISPR arrays present within and across samples of the same individual. A CRISPR array was considered nested within another CRISPR array if both shared the same or highly similar repeats (>90% identity and coverage, clustered using CD-HIT-Est with option -*g* 1) and all the spacers of the former were also harbored by the latter. Group-representative arrays were identified as those that were not nested within any other array. Finally, clusters of CRISPR arrays were built by considering all the arrays nested within the same group representative. This algorithm allowed us to retrieve clusters of highly similar CRISPR arrays while accounting for incomplete assemblies and spacer gain and loss over time. Because of that, we refer to these clusters as CRISPR lineages or lineages of arrays. Note that, although the strict nesting criterion can leave partially overlapping arrays unmerged, this minimizes the risk of generating heterogeneous clusters with lineages that share conserved spacers but behave independently in the timescale of the study.

### Search for CRISPR targets in the local microbiome

Protospacers in the local microbiome were identified by running nucleotide BLAST against metagenomic contigs previously assembled by HMP2-IBD, with spacers as queries, options “-dust no -word_size 8”, and a 95% identity and coverage thresholds. We discarded hits to known CRISPR arrays (identified with CRISPRCasTyper) and hits located within a 100bp neighborhood of known CRISPR repeats (found with nucleotide BLAST, options “-dust no - word_size 8”, 95% identity and coverage).

One spacer in the dataset produced an extremely high number of hits to the local metagenome (16,327 hits, while the second largest number was 121). Closer inspection revealed that the spacer was highly similar to a transposase sequence found in bacterial chromosomes and prophages. To avoid biases in downstream analyses, the spacer was removed from the dataset.

### Identification of spacer targets

The targets of CRISPR of spacers were inferred by performing a nucleotide BLAST search of all spacers against a merged database of mobile genetic elements (MGEdb), setting BLAST parameters for short queries (-dust no -word_size 8) and keeping all hits with >95% identity and coverage. The merged MGEdb included 198,607 complete viral genomes from the NCBI Genome database (downloaded on 27/01/2021 from ftp.ncbi.nlm.nih.gov/genomes/Viruses/) and the NCBI Virus database (downloaded in interactive mode on 27/01/2021 from https://www.ncbi.nlm.nih.gov/labs/virus/vssi/%23/virus?SeqType_s=Nucleotide&Completeness_ s=complete), a set of 1,887 gut phages recently identified by Benler et al. (56), a set of 13,274 ICE/IME from Botelho (57), and all sequences from mMGE v1.0 (58) (494,835 phages and 92,493 plasmids), ImmeDb (59) (201 phages and 4,727 ICE/IME, transposable elements, and other unclassified MGE), ICEberg3 (60) (4,461 ICE/IME and other chromosomally integrated elements), and IMG/PR (61) (699,973 plasmids). Based on the BLAST results, we classified spacers according to their most likely target (virus, plasmid, or other mobile genetic elements). Spacers with inconsistent target predictions were classified as “ambiguous”. Spacer-target assignments were annotated according to the accuracy of the match as identical (100% identity and coverage) or highly similar (identity and coverage above 95% but below 100%). Moreover, viral targets were taxonomically annotated according to the information in MGEdb. Spacers targeting phages from the infant gut were identified by running BLAST searches against the database generated by Shah et al (62).

To infer the target of spacers that did not match any sequence in MGEdb but had protospacers in the local microbiome, we extracted up to 15 kb neighborhoods (7.5 kb upstream and downstream from the protospacer, which corresponds to the mean length of phage genomes in MGEdb) from the HMP2-IBD contigs and ran a nucleotide BLAST search (option dc-megablast) against MGEdb setting an e-value threshold of 10^-8^. If the search returned hits from multiple MGE, we discarded those with bitscores lower than 0.9 times the bitscore of the best hit. Compared to just keeping the best hit, this approach preserves the uncertainty among high-scoring candidates. Spacers with inconsistent predictions were classified as “ambiguous” and those whose target remained unclassified after this second step were labeled as “unknown”.

### Prediction of lytic or lysogenic phage lifestyle

Phages targeted by CRISPR spacers were classified as lytic or lysogenic based on the presence of integrases in phage genomes. To that end, we extracted 49,645 phage genomes from MGEdb and 110,712 metagenome-assembled contigs from HMP2-IBD that contained protospacers and had been previously characterized as viral (see previous section). Then, we ran Prodigal v2.6.3 (63) with the option *-p meta* to identify open reading frames and searched for hits to viral integrases using the *hmmersearch* command of the HMMER v3.3.2 suite. We included the following PFAM profiles for integrases: PF00239, PF00589, PF02899, PF07508, PF12482, PF12834, PF13009, PF13102, PF13495, PF14659, PF16795, and PF18644. The search returned 11,631 viral sequences with putative integrases (e-value <10^-8^) that also contained protospacers. The corresponding spacers (27,868 in total) were regarded as targeting lysogenic phages.

### Prevalence and abundance of CRISPR targets in the local microbiome

To study the quantitative dynamics of CRISPR targets in the microbiome, we focused on spacer-protospacer pairs present in samples from the same individual (possibly at different times) that fulfil the following conditions: (1) the first appearance of the spacer occurs later than the first appearance of the CRISPR lineage with which the spacer is associated; and (2) spacers are present in at least 2 samples. These two conditions allowed us to compare target abundances before and after the first appearance of the spacer, while controlling for the intermittent prevalence of hosts (using CRISPR lineages as proxies for the latter). For the same reason, we excluded from these analyses all samples in which the CRISPR lineage was not detected. After applying these filters, 1540 spacer-protospacer pairs remained, representing 1057 CRISPR lineages and 5369 protospacer occurrences.

To estimate the prevalence of targets, we counted the number of samples that contain the protospacer before, at, and after the first appearance of the spacer, pooled these counts for all spacer-protospacer pairs, and normalized them by the number of samples in each group. Confidence intervals were calculated using the Clopper-Pearson method as implemented in the *statsmodels* Python library. To quantify target abundances, we extracted 4kb neighborhoods (2kb upstream and downstream) of protospacers from HMP2-IBD contigs and mapped metagenomic reads to those neighborhoods with CoverM v0.6.1. The relatively small size of these neighborhoods was chosen to minimize the risk of including flanking regions from host genomes in the case of integrated MGE. Sample-wise relative abundances were calculated as the number of mapped reads divided by the length of the mapped sequence (generally 4kb, except for protospacers located near contig ends) and by the total number of reads in the sample. The sample-wise abundance in samples in which the target was not detected was set to 0. Finally, to facilitate the comparison of trends across targets, we normalized the sample-wise abundances of each target by the maximum abundance observed for that target in the whole time series.

### Taxonomic assignation of CRISPR-Cas systems

To assign a taxonomic origin to CRISPR arrays, we leveraged the fact that CRISPR repeats are taxon-specific (although the taxonomic level of such specificity varies among repeats) (64). First, we ran CRISPRCasTyper on the preassembled HMP2-IBD metagenomic contigs to predict CRISPR arrays. Then, we used CD-HIT to identify contigs from HMP2-IBD that contain CRISPR arrays with the same repeats (90% identity and coverage) as those assembled by CasCollect. Those contigs were provided as inputs to the program CAT/BAT v5.2.3 (65) for taxonomical assignment at the lowest non-ambiguous available level. For CRISPR arrays whose repeats could not be mapped to any HMP2-IBD contig, we directly ran CAT/BAT on the contigs obtained with CasCollect. (Note, however, that the level of resolution obtained with CasCollect contigs was generally lower due to their shorter length.) Finally, all CRISPR arrays that remained unassigned and were part of a lineage with one or more assigned members were given the same taxonomical origin as the other members of the lineage, as long as no inconsistencies were found.

### Spacer location bias and asymmetry

We studied the relative position of spacers that target phages from the local metagenome (henceforth “locally adapted spacers”) in a subset of 4,054 oriented arrays with >5 spacers, separately considering identical and highly similar hits (CRISPR-Cas subtypes with <25 valid arrays were not included in this analysis). To avoid artifacts associated with finite-size effects in small arrays, we used a sliding-window approach to estimate the density of locally adapted spacers along the array. We set a window width equal to 12% of the total length of the array (equivalent to 2/3 of the inverse of the median number of spacers) and a step equal to 1.5% of the total length of the array (1/8 of the window width). This choice results in a window size slightly smaller than the typical span of a spacer (given by the inverse of the median number of spacers per array), which provides a good balance between resolution and smoothness.

### Quantification of transcription levels along CRISPR arrays

For 753 samples from HMP2-IBD with available metatranscriptomes, we mapped metatranscriptomic reads to CRISPR assemblies obtained from the same samples with minimap2 v2.24, setting the option ‘-x sr’ for short reads (66). To improve specificity towards spacers in CRISPR arrays, we also included contigs with protospacers in the reference genomic dataset and removed all mapped reads with aligned lengths below 70 nucleotides using the command FilterSamReads from gatk 4.1.2.0 (67). This 70-nucleotide threshold was chosen to ensure that alignments covered the equivalent of one spacer and one repeat (the average spacer and repeat lengths in our dataset are 34 and 31, respectively). The coverage per position was calculated with the command genomeCoverageBed from BEDTools 2.28.0 (68).

### Clustering and classification of CRISPR-spacer-target dynamics

To study the temporal dynamics of CRISPR lineages, spacers, and targets, we first identified all the spacers that were observed for the first time later than their corresponding CRISPR lineage. Besides this basic requirement, we did not impose any further condition to filter potential acquisition events (for example, requiring that the new spacer was located at the leading end) because predicted orientations were only available for a small fraction of arrays. (Moreover, we could not discard that more than 1 spacers were incorporated between two sampling points, which would make newly acquired spacers look as if they were internal.) We codified the time series associated with each spacer as a string of characters, using letters A-F to indicate all possible combinations for the presence or absence of that particular spacer, its target, and its corresponding CRISPR lineage. “Empty” states (that is, those characterized by the absence of spacer, CRISPR lineage, and target) were removed from both ends of the string and collapsed to a single instance if appearing more than once between non-empty states (see Supplementary Note S1 for a detailed justification of this procedure). To cluster similar time series and obtain representative trajectories, we performed pairwise alignments of all strings with the Biopython command pairwise2.align.globalms (69) using the following scores: 1 per match, -2 per mismatch, -2 per gap opening, and -1 per gap extension (see Supplementary Note S1 and Figure S3 for further details and discussion of alternative scoring schemes). After normalizing the scores by the length of the longest member of each pair and discarding pairs with negative scores, we built a time-series similarity network using normalized scores as weights. Time series were then clustered with Infomap v0.21.0 (70), with options -2 -u -N 200 (two-level partition assuming undirected links and selecting the best of 200 trials).

## Results and discussion

### Assembly and characterization of CRISPR arrays in the human gut microbiome

We combined state-of-the-art methods to assemble CRISPR arrays from the gut microbiomes of 130 individuals, recurrently sampled every 2 weeks over a 1-year period as part of the integrative Human Microbiome Project (Figure 1a). After curation and removal of low-quality arrays, we obtained a set of 79,475 CRISPR arrays, encompassing 11,373 unique repeats and 153,745 unique spacers. Of those, around 4,300 arrays were adjacent to a Cas operon, what allowed us to predict the direction of transcription by applying three alternative criteria (see Methods). We identified CRISPR arrays in 36 out of the 40 most abundant bacterial genera in all gut samples, which collectively account for 94% of bacterial abundance (Suppl. Table S2). The number of unique CRISPR arrays per sample broadly varies through time and across microbiomes, with significantly lower numbers in microbiomes obtained from patients with Crohn’s disease and ulcerative colitis (Figure 1b-c). Such lower numbers are consistent with the reduced microbial diversity associated with dysbiosis in inflammatory bowel diseases (44).

**Figure 1:**
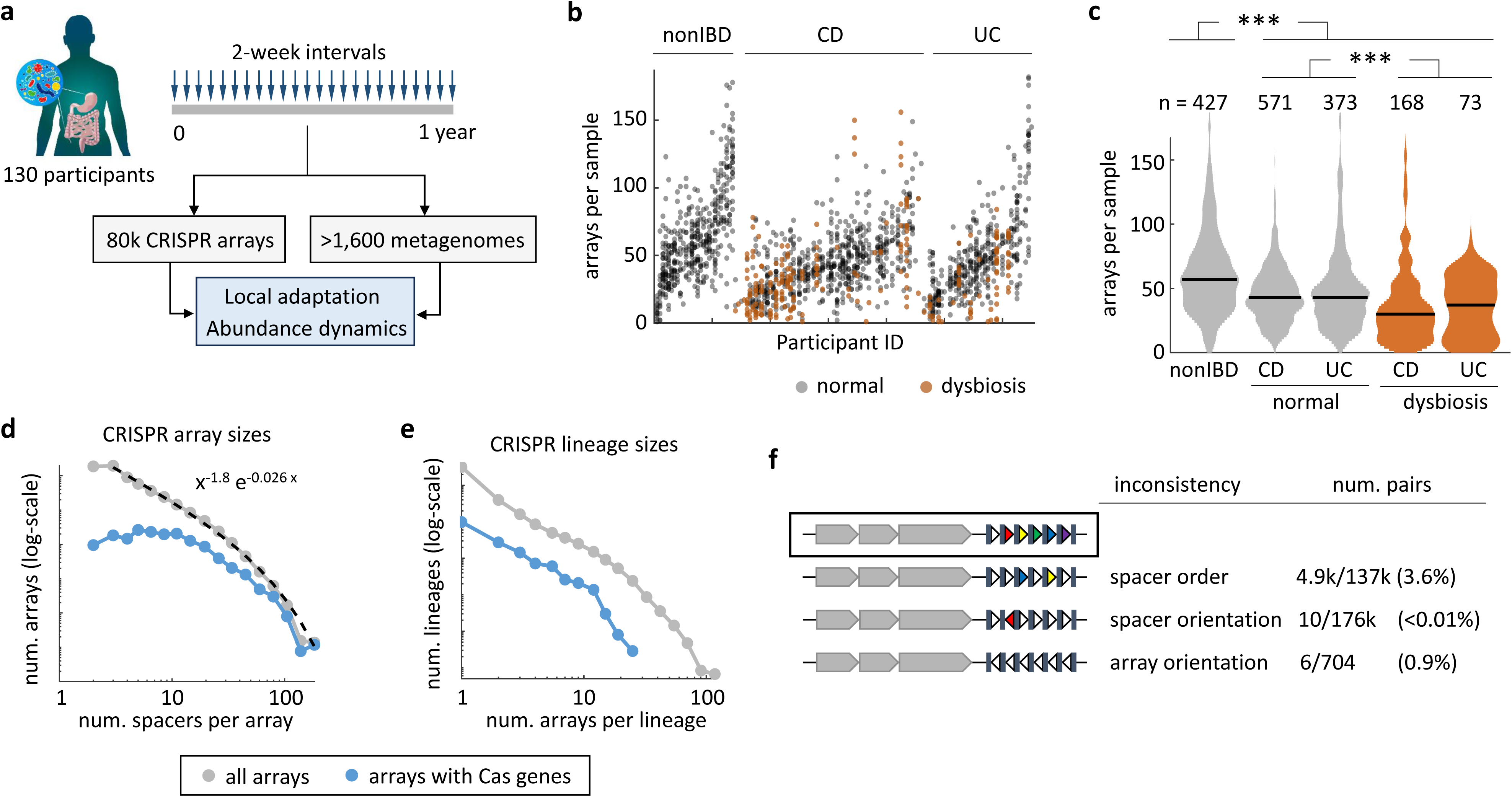
Assembly and quality-control of CRISPR arrays in the human gut microbiome. (a) Schematics of the study design based on metagenomic data from the HMP2-IBD cohort. CRISPR arrays were assembled de novo with CasCollect. (b) Number of CRISPR arrays per sample. (c) Distribution of the number of CRISPR arrays per sample in patients with Crohn’s disease (CD), ulcerative colitis (UC), and participants with no inflammatory bowel disease (nonIBD). The central lines indicate the medians for each group. Kruskal-Wallis omnibus test for differences among groups (χ^2^ = 148; *p* < 10^-10^). Mann-Whitney tests for differences between nonIBD and disease (Z = 10.4; *p* < 10^-10^); and between normal and dysbiosis (Z = 6.76; *p* < 10^-10^). (d) Distribution of the number of spacers per CRISPR array, in all arrays (grey) and arrays located in the vicinity of Cas genes (blue). The dashed line indicates the best fit to a truncated power-law. (e) Distribution of the number of CRISPR arrays per lineage (note the logarithmic scale in the y-axis, also in (d)). (f) Quantification of the number of pairs of CRISPR arrays from the same lineage that show inconsistencies in spacer order, spacer orientation, and array orientation.

The CRISPR array sizes follow a heavy-tailed distribution that is well-fit by a truncated power law with exponent close to -2 and a cutoff at around 100 spacers (Figure 1d). Similar distributions, although with higher cutoffs, have been reported for CRISPR arrays from complete genomes (71, 72). Thus, the high heterogeneity of array lengths observed in this dataset likely reflects a true property of CRISPR systems in the gut microbiome, rather than an excess of fragmentary or poorly-assembled arrays.

We applied a strict nesting criterion to group CRISPR arrays into 41,742 clusters, of which 9,870 contain >1 array (Figure 1e). These clusters represent lineages of arrays from the same microbiome (possibly sampled at different times) that are likely related by sequential gain and loss of spacers. Array lineages serve a triple purpose in this study. First, lineages are pivotal to assess the consistency of CRISPR assemblies by comparing spacer order and orientation in arrays from the same lineage. Second, lineages permit a robust characterization of CRISPR diversity by minimizing the impact of incomplete arrays and abundance biases. Third, lineages constitute a convenient proxy for the presence of subpopulations of hosts in the microbial community.

Within-lineage comparisons show that spacer order and orientation are highly consistent across pairs of related arrays (Figure 1f). Such high consistencies are not a by-product of the method used to group arrays into lineages, which was solely based on spacer presence or absence and did not discriminate between spacers and their reverse complements. Overall, internal rearrangements only affect 3.6% of all array pairs, whereas spacer inversions are virtually inexistent (<0.01%). Predicted array orientations are also generally consistent with respect to spacer order, with inconsistencies observed in <1% of the pairs. Because lineages include arrays separately assembled in different samples, within-lineage consistency strongly suggests that CRISPR assemblies are reliable in terms of spacer order and orientation.

The most abundant CRISPR-Cas systems in the human gut microbiome belong to subtypes I-B, I-C, I-E, and II-C (Figure 2). Subtypes I-B, I-C, and II-C are most often associated with Firmicutes and Bacteroidetes, whereas subtype I-E is observed in Firmicutes and Proteobacteria. Alterations in the composition of the microbiome significantly affect the profile of CRISPR subtypes in Crohn’s disease and ulcerative colitis (Figure 2 and Suppl. Table S2). In agreement with previous studies (44, 73), dysbiosis leads to a decrease in the relative proportion of subtypes I-B, I-C, I-E, and II-C that can be attributed to the depletion of *Faecalibacterium prausnitzii* (I-C and I-E) and several groups of Bacteroidetes (I-B, I-C, and II-C). Among the less abundant subtypes, IV-A (typically carried by large conjugative plasmids (74)) stands out for its strong association with normal (that is, no dysbiotic) samples.

**Figure 2:**
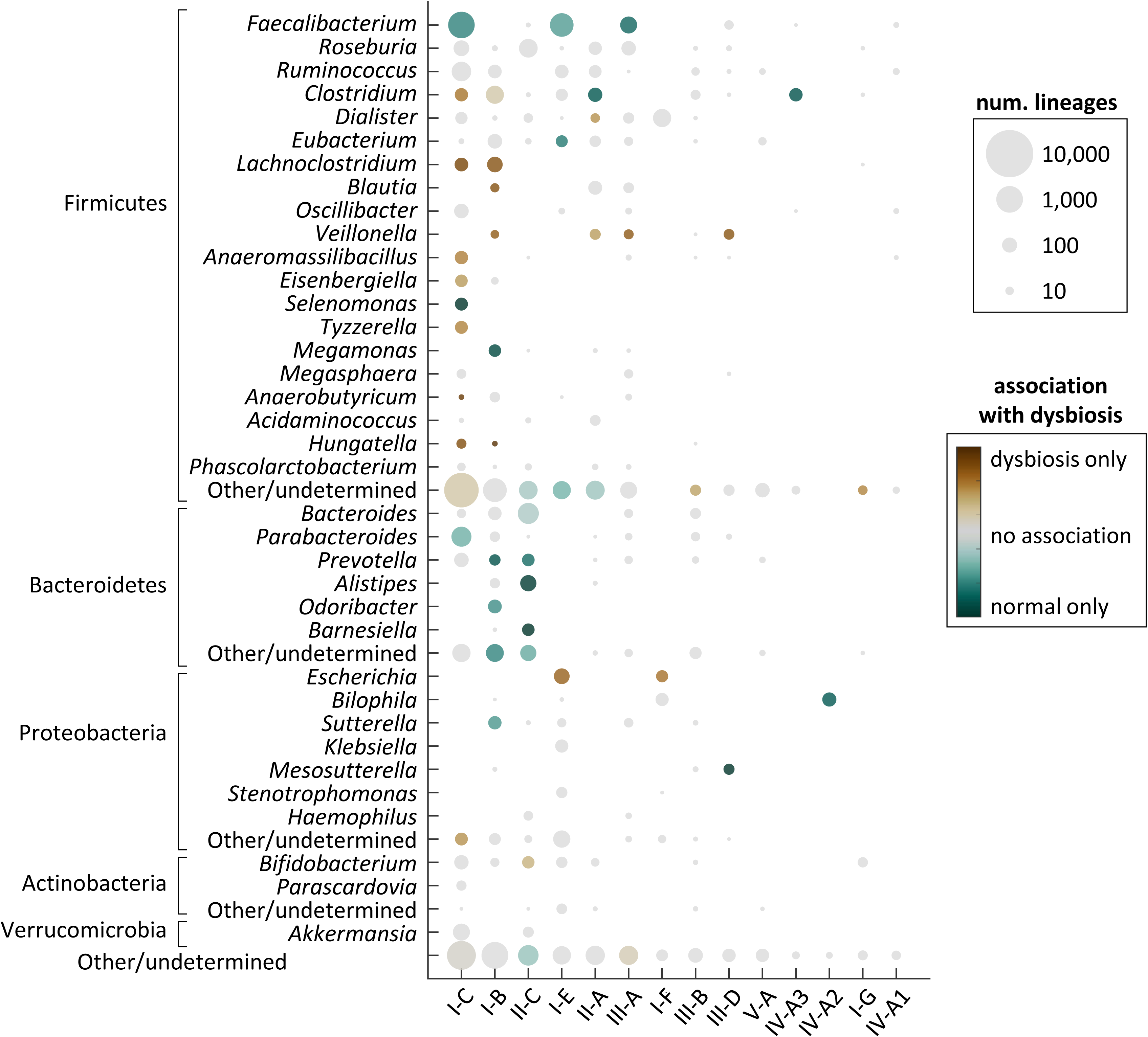
Distribution of CRISPR subtypes among taxa and association with dysbiosis. The area of each circle is proportional to the number of CRISPR lineages in the whole dataset. Colors indicate the relative association with dysbiosis. Subtypes with <50 CRISPR lineages are not represented. “Other/undetermined” includes genera (or phyla) with <50 CRISPR lineages and lineages with ambiguous classification at the genus (or phylum) level. CRISPR subtypes on the x-axis are listed from left to right based on their total abundance.

### Selection promotes long-term immunity against endemic MGE

To identify the potential targets of CRISPR immunity, we mapped the spacers in CRISPR arrays to a large collection of phages, plasmids, and other MGE obtained from state-of-the-art (meta)genomic databases (see Methods). Overall, 41% of the spacers show high similarity (zero or one mismatch) to known phages and only 4% map to plasmids and other MGE (Figure 3a). These values are notably higher than those reported by global surveys of CRISPR spacers from complete prokaryotic genomes (75, 76) but close to those obtained for gut bacteria (43, 77), probably reflecting the higher coverage of MGE from the human gut in public databases. Around 6% of the spacers target sequences from the local microbiome (defined as the union of all samples collected from the same individual). We refer to these spacers as “locally adapted” because they can provide immunity against MGE that are locally observed in the 1-year period covered by the dataset. The most frequent targets of locally adapted spacers are lysogenic phages (43%), followed by lytic and other non-lysogenic phages (28%), plasmids (16%), and other MGE (1%). Around 10% of the spacers (labeled as “ambiguous” in Figure 3a-b) match sequences found in multiple classes of MGE. All in all, we could assign a unique or ambiguous origin to 99% of all locally adapted spacers. Therefore, the disproportionate number of hits to phages and plasmids compared to other MGE is probably genuine, despite the admittedly biased representation of different classes of MGE in public databases. Comparing among CRISPR subtypes (Figure 3b), local adaptation is more frequent in subtypes I-C and II-C (7.7-8.5% of spacers are locally adapted, compared to 2.5-4.2% in other subtypes; χ^2^ test *p* < 10^-10^). Moreover, subtypes II-A, II-C, and III-A target a higher proportion of plasmids than class I systems (12% in I-C vs 25% in II-C; Fisher’s exact test *p* < 10^-10^).

**Figure 3:**
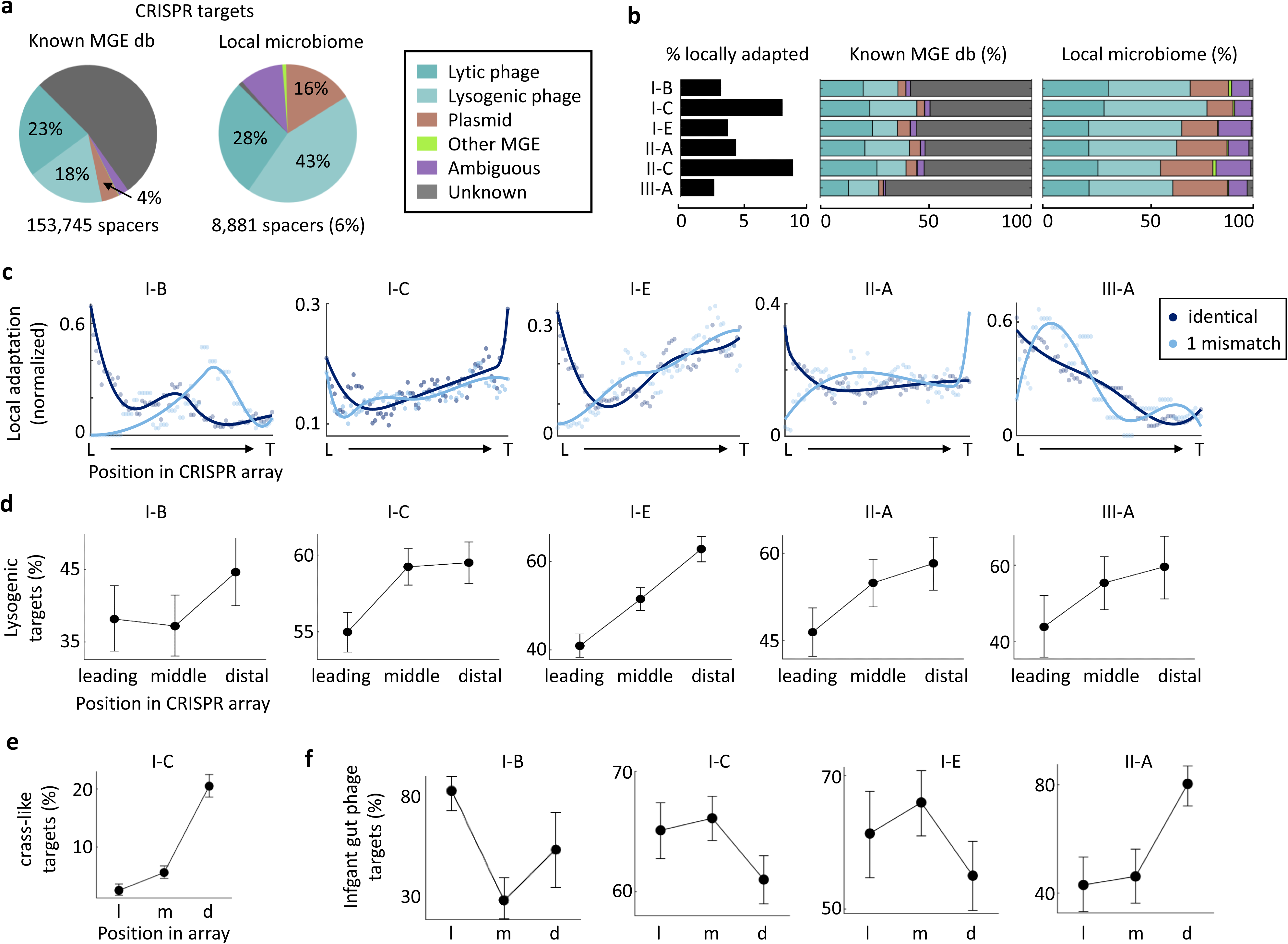
Targets of CRISPR-Cas immunity in the gut microbiome. (a) Relative representation of different classes of mobile genetic elements among the targets of CRISPR-Cas systems. Left pie chart: hits to a merged database of known mobile genetic elements (MGE db). Right: protospacers from the local microbiome, defined as the combination of all samples obtained from the same participant. The segment for lytic viruses also includes chronic (non-lysogenic) viruses (b) Distribution and functional profile of CRISPR targets across CRISPR-Cas subtypes. Locally adapted spacers (bar plot on the left) are those that target sequences from the local microbiome. (c) Relative position of locally adapted spacers within the CRISPR array. L: leading end; T: trailing end. (d-f) Fraction of spacers targeting lysogenic (d), crass-like (e), and infant gut (f) phages (with respect to all spacers that target phages) in the leading, middle, and distal sections of the array. Whiskers represent 95% confidence intervals calculated with the Clopper-Pearson method. In (b) and (c), only those subtypes with >100 locally adapted spacers are shown. Panels (d-f), only include subtypes with >100 spacers targeting lysogenic phages (d), crass-like phages (e), and infant gut phages (f) within high-quality, locally adapted arrays (oriented and with ≥5 spacers). Despite the high fraction of locally adapted spacers, subtype II- C was excluded from (c-f) due to the unavailability of accurate predictions for array orientation.

Locally adapted spacers are not uniformly distributed along the CRISPR array (Figure 3c). On the contrary, they tend to concentrate near both ends of the array. The enrichment at the leader (or proximal) end is consistent with recent spacer acquisitions targeting MGE from the local microbiome. However, the excess of locally adapted spacers near the trailer (or distal) end, especially evident in subtypes I-C and I-E, is somewhat paradoxical. In fact, distal spacers are generally the oldest in the array and their role in immunity is a matter of debate (78, 79). Three lines of evidence reject the possibility that locally adapted spacers near the distal end represent a technical artifact. First, array orientation and spacer order are highly consistent across arrays from the same lineage, despite the arrays being independently assembled from different samples. Second, the same trend is observed when the orientation of the arrays is inferred with three independent methods (Cas Orientation, CRISPRDirection, and CRISPRLoci; Figure S2). Third, in most subtypes, mismatches between the spacer and target sequences become more frequent towards the trailer end, as expected for correctly assembled arrays. CRISPR subtypes I-C and I-E contain 85% of all locally adapted spacers within the set of high-quality arrays (those with adjacent Cas genes and 5 or more spacers). Therefore, despite the concentration of locally adapted spacers near the trailer end is not observed in all subtypes, it arguably represents a dominant trend in the gut microbiome.

We conceive two possible biological explanations for the observed trend in the distribution of locally adapted spacers. The first is that CRISPR-based immunity leads to the complete elimination of MGE targeted by spacers in the middle section of the array, whereas MGE targeted by distal, less efficient spacers can persist in the microbiome. In this scenario, most middle spacers would not be identified as “locally adapted” simply because they succeeded in eliminating their targets. Although CRISPR immunity can effectively prevent MGE proliferation in culture (80, 81), it is less likely that complete elimination of MGE occurs in complex, spatially extended microbial communities (26, 82) (and the human gut virome appears indeed highly stable over time (83)). More generally, this scenario does not account for the possibility that the same MGE is targeted by multiple spacers located at different positions in different arrays, which would effectively decouple within-array spacer positions and population-level MGE abundances.

The second (and more likely) explanation is that locally adapted spacers near the trailing end are a consequence of sustained selection for long-term immunity against endemic MGE. It is widely accepted that intense selection and frequent spacer loss via homologous recombination shape the composition of CRISPR arrays in natural populations, promoting the maintenance of functional spacers (42). Therefore, if endemic MGE impose a sustained burden on the population, it can be expected that older spacers providing with effective long-term immunity against such MGE accumulate at the distal part of the array. This phenomenon has been observed in *Leptospirillum* strains from an acid mine drainage where the same phage population persisted for at least 5 years (23, 84). We propose a similar explanation in the case of the adult human gut, where bacteria coexist with a very stable virome dominated by crass-like phages (83, 85–87) and are exposed to continuous (but low-intensity) induction of prophages (88, 89). In support of the long-term memory role of distal CRISPR spacers, we found that the fraction of spacers targeting lysogenic and crass-like phages systematically increases from the leading to the trailing end of the array (Figure 3d-e).

Considering the high rates of lytic activity and CRISPR spacer turnover that characterize the infant gut, and the possible contribution of early phage-bacteria interactions to shaping the adult gut microbiome (13, 62, 90, 91), an intriguing possibility might be that long-term memory spacers were selected during the consolidation of a coexisting community of MGE and bacteria in the first years of age. To test that hypothesis, we investigated the relative abundance of spacers targeting phages from the infant gut in leading, middle, and trailing regions of CRISPR arrays (Figure 3f). Interestingly, spacers that target infant gut phages do not concentrate at the trailing end (except in subtype II-A), but they represent 60-80% of all the phage-targeting spacers in the leading region of the array. These results confirm that exposure to phages in the infant gut is a major trigger of CRISPR adaptation, though it does not seem to affect selection for long-term immune memory.

To provide efficient long-term immune memory, spacers near the trailing end of the array must be expressed at sufficient levels to form effector complexes. Indeed, lack of expression due to early termination of transcription limits the functionality of distal spacers in experimental conditions (39–41). By analyzing metatranscriptomes, we found that distal spacers are moderately expressed, with transcript abundances around 30-40% of those observed for spacers near the leading end (Figure S4). Distal spacers may also contribute to long-term immunity through primed adaptation, that is, by promoting the acquisition of additional spacers from cognate MGE (92). Primed adaptation is widespread in type I and II CRISPR systems (93, 94) and operates for spacers that do not exactly match their target (95) or are expressed at low levels (96). Because spacers acquired through this mechanism are often biased in terms of genomic location and strand specificity, indirect evidence for primed adaptation can be obtained by testing for non-random patterns in the relative location of protospacers within the target genome. Following that strategy, we identified several instances of MGE targeted by >1 spacer from the same array, which could correspond to primed adaptation. However, the number of instances observed was insufficient for statistical confirmation.

### CRISPR-Cas immunity modulates microbiome composition

The longitudinal nature of the HMP2-IBD metagenomic survey allowed us to investigate how CRISPR immunity affects the dynamics of MGE and bacterial hosts in the gut microbiome. We first studied if the appearance of a new spacer in a preexisting CRISPR lineage (possibly implying a spacer acquisition event) was associated with changes in the prevalence and abundance of the target MGE (Figures 4a and S5a). Consistent with the expectation that exposure to MGE induces immune adaptation, we found that putative spacer acquisitions occur in samples in which the abundance of the target MGE is higher than average. In sharp contrast, subsequent observations of the spacer are associated with significantly decreased MGE abundances. Notably, such decrease is not observed in populations that have seemingly lost the spacer but in which the CRISPR lineage (and therefore the host) is still detected. As expected, these trends are strongest in the case of lytic and lysogenic phages, but also significant for plasmids. Despite its quantitative effect on MGE abundance, CRISPR-Cas immunity does not generally lead to the complete elimination of its targets. Indeed, around 50% of lytic and lysogenic phages targeted by CRISPR-Cas can still be detected following the acquisition of a cognate spacer in the population (Figures 4b and S5b). In the case of plasmids, acquisition of cognate spacers does not produce a significant drop in prevalence. All in all, these observations imply that CRISPR-Cas immunity modulates phage and plasmid abundances in the human gut, while not compromising their long-term persistence. This conclusion is consistent with previous observations from free-living communities and with the theoretical prediction that CRISPR-Cas immunity can stabilize complex phage-host populations through eco-evolutionary feedback (28, 97–99).

**Figure 4:**
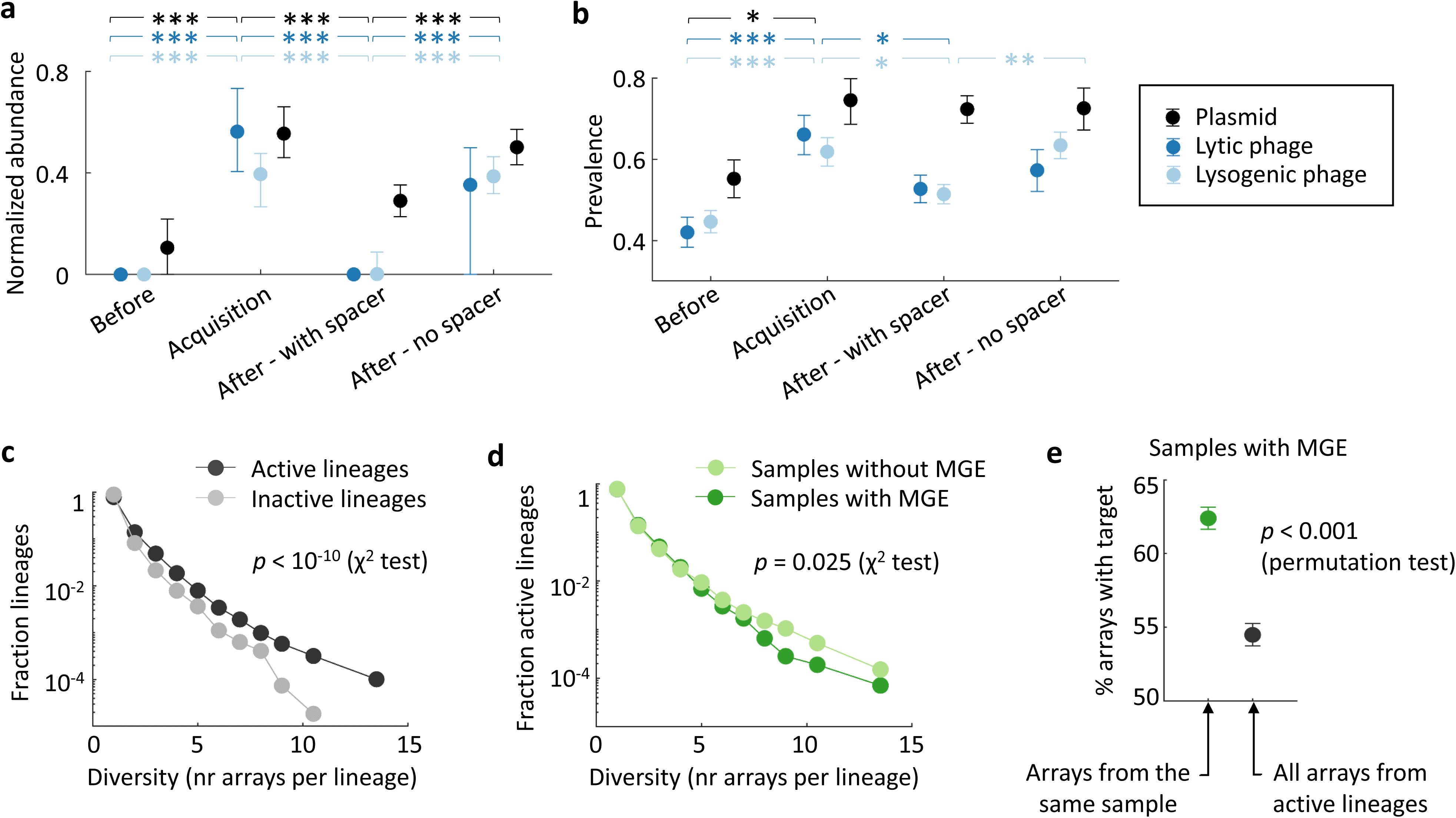
Effect of CRISPR-Cas immunity on MGE abundance and host diversity (see also Figure S5). (a) Median normalized abundance of targeted MGE (plasmids, lytic/non-lysogenic phages, and lysogenic phages) before the spacer was first observed in the host population (“Before”), in the sample in which the spacer was observed for the first time (“Acquisition”), in subsequent samples in which the spacer was present (“After – With spacer”), and in subsequent samples that contained the same CRISPR lineage but not the spacer (“After – No spacer”). Error bars indicate the 95% confidence intervals for the median, based on 2,000 bootstraps. All comparisons shown above the plot are statistically significant with *p* < 10^-4^ (Mann-Whitney test). Note that comparisons are conservative with respect to array incompleteness, because apparent loss of spacers in incomplete arrays would lead to misclassification of samples (from “After – With spacer” to “After – No spacer”), therefore reducing the observed differences. (b) Prevalence of targeted MGE, calculated as the fraction of samples in which the target is detected. Error bars indicate the 95% confidence intervals (Clopper-Pearson method). Statistical significance based on Fisher’s exact test: *p* < 0.05 (*), *p* < 0.01 (**), *p* < 0.001 (***). (c) Diversity distribution (number of distinct arrays) of CRISPR lineages that target at least one MGE from the local microbiome (“active” lineages) and those that do not show any sign of local adaptation (“inactive” lineages). (d) Number of arrays per lineage in the presence and absence of MGE. For a given active lineage, samples with MGE are those that contain at least one MGE targeted by at least one spacer from that lineage. (e) Percentage of arrays per lineage that target MGE from the same sample, compared to the expected value for randomly chosen arrays. The randomized set only includes active lineages and samples with MGE targeted by those lineages. Error bars indicate the 95% confidence intervals based on 2,000 bootstraps.

We then investigated if the interplay of CRISPR immunity and phage predation affects host diversity at the population level. To that end, we divided CRISPR lineages into those that are possibly involved in effective immune interactions (by harboring at least one locally adapted spacer) and those that do not show any evidence of local adaptation. The former tend to be more diverse, that is, they encompass more array variants per lineage, likely reflecting higher rates of spacer acquisition and loss (Figure 4c). Moreover, the diversity of CRISPR arrays in these lineages is lower in samples in which targeted phages are present (Figure 4d). This observation underscores the occurrence of weak MGE-induced selective sweeps in the host population, that transiently favor those hosts bearing CRISPR arrays with cognate spacers (Figure 4e). The weakness of these selective sweeps could in part be explained by clonal interference among multiple immune lineages, as previously suggested for hot spring communities (27), and by the spatially structured nature of the gut microbiome.

### Temporal dynamics of MGE and CRISPR arrays

To better understand the interplay between CRISPR-Cas immunity and the dynamics of MGE and hosts, we selected 1,769 time series for single spacer-target pairs whose dynamics are compatible with possible spacer acquisition events at the population level. Then, we applied an unsupervised clustering algorithm to group similar trajectories based on the temporal patterns of presence or absence of spacers, targets, and CRISPR lineages (the latter serving as proxies for host subpopulations). Further details about this procedure are provided in the Methods section, Supplementary Note S1, and Figure S3, where we show that the final clusters are stable with respect to small methodological variations. Despite the potentially combinatorial diversity of individual trajectories, we found that 40% of them belonged to one of the 10 largest clusters. The inspection of such representative clusters reveals qualitative differences across MGE in their temporal dynamics (Figures 5 and S6). Phages are preferentially associated with episodic dynamics (TC-1, 3, 5, and 7) involving type I-C CRISPR-Cas systems. In turn, plasmids display recurrency (TC-13 and 14) and are most often targeted by type II-C systems. Although array incompleteness could, in principle, introduce artifacts in time series in the form of apparent spacer loss with lineage persistence, such transitions were rare or absent from the major trajectory clusters (Figure 5a).

**Figure 5:**
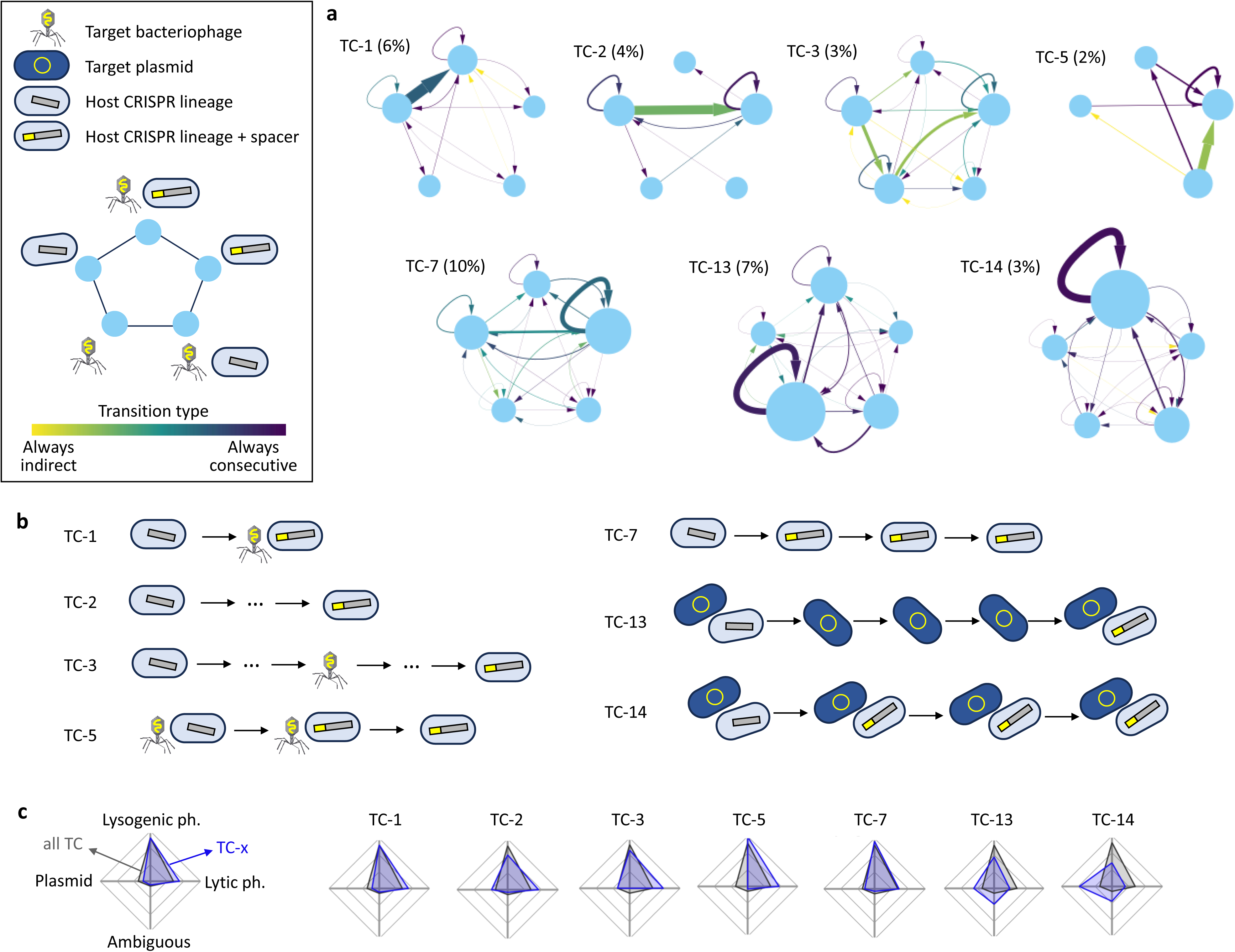
Temporal dynamics of MGE and CRISPR arrays. (a) Network representation of 7 of the most frequent trajectory clusters (TC) representing potential immune adaptation in the host population. The percentages in parentheses indicate the relative frequency of each trajectory. Each node represents a state, defined by the presence or absence of a spacer, its cognate MGE, and the CRISPR lineage in which the spacer was acquired (legend in the upper left box). Node sizes are proportional to the number of times that a state is observed in a trajectory. Arrow widths represent transition probabilities between states and the color scale indicates the fraction of direct vs indirect transitions (direct transitions correspond to two consecutive samples, indirect transitions involve intermediate samples in which neither the MGE nor the CRISPR lineage were observed). (b) Simplified representation of typical trajectories associated with the TC in (a). (c) Relative association of lytic (or non-lysogenic) phages, lysogenic phages, plasmids, and ambiguous MGE with each TC.

One of the most frequent trajectories in the data set (TC-1) corresponds to a single observation of a CRISPR lineage followed by a simultaneous observation of a new spacer and the cognate phage, in what might represent a spacer acquisition event. Other representative trajectories involve variations of the same theme, with loss of the phage following spacer acquisition (TC-5) or missing the intermediate acquisition event (TC-2 and 3). In contrast, the most frequent trajectories associated with plasmids represent two alternative scenarios, one dominated by the plasmid (TC-13) and the other characterized by the stable coexistence of plasmid-bearing and plasmid-resistant host subpopulations (TC-14). The latter is reminiscent of the global population structure of *Pseudomonas aeruginosa* , in which spacers from active CRISPR-Cas systems target MGE integrated in closely related genomes that lack CRISPR (100), and in hot spring populations of *Hydrogenobacter sp* ., where the spacers of single-cell assembled genomes target MGE found in other cells from the same population (101). Our results suggest that such crosstalk between CRISPR-Cas systems and plasmids that coexist in the host population is also widespread in the human gut.

## Conclusions

By analyzing a large metagenomic dataset encompassing 130 individual gut microbiomes periodically sampled for up to 1 year, we found that locally adapted spacers targeting lysogenic and crass-like phages are often located in the distal region of CRISPR arrays. This finding strongly suggests that selection promotes the maintenance of long-term memory spacers against endemic MGE, and that periodic reactivation of prophages is an important stressor for the human gut microbiome. Interestingly, leader and trailer regions of CRISPR arrays appear to be specialized in short- and long-term memory, respectively. We expect that selection for long-lasting immunity and polar allocation of long-term memory spacers are not unique to the human gut microbiome, but general features of microbial communities with stable viromes. Testing such prediction constitutes an interesting subject of future research.

A second major finding of this study was that the interaction of phages and CRISPR-Cas immunity quantitatively affects the composition of the human gut microbiome. Phage and plasmid abundances tend to decrease after the first appearance of cognate CRISPR spacers in the microbiome, although such decrease does not generally lead to their complete elimination. On the other side, in the presence of MGE, bacterial populations experience weak and incomplete selective sweeps favoring immune lineages. The coexistence of phages and diverse host lineages despite effective CRISPR-Cas immunity suggests that spatial heterogeneity, clonal interference, and fluctuating selection could have a major role in explaining phage-bacteria interaction dynamics in the gut. Disentangling the contribution of these mechanisms warrants further investigation, as they may differentially affect the efficacy of phage therapy and other intervention strategies aimed at controlling the gut microbiome.

## Acknowledgements

A.L-B. is supported by the Agencia Estatal de Investigación of Spain (Grant No. PRE2020-092935). J.B. is supported by the Maria Zambrano grant of the Spanish Ministry of Universities (Grant No. UP2021-035), and the Severo Ochoa Program for Centres of Excellence in R&D of the Agencia Estatal de Investigación of Spain (Grant No. CEX2020-000999-S (2022–2025) to the CBGP). J.I is supported by the Ramón y Cajal Programme of the Spanish Ministry of Science (Grant No. RYC-2017–22524); the Agencia Estatal de Investigación of Spain (Grant No. PID2019-106618GA-I00), the Severo Ochoa Programme for Centres of Excellence in R&D of the Agencia Estatal de Investigación of Spain (Grant No. SEV-2016–0672 (2017–2021) to the CBGP); and the Comunidad de Madrid (through the call Research Grants for Young Investigators from Universidad Politécnica de Madrid, Grant No. M190020074JIIS).

## Data availability

All relevant data produced in this study have been deposited in the Zenodo repository (DOI: 10.5281/zenodo.10555110) and are freely available.

