## Supplementary Notes, Figures and Table S1 for "Dynamics of CRISPR-mediated virus-host interactions in the human gut microbiome"

This document contains Supplementary Note S1, Figures S1-S6, and Table S1. Supplementary Table S2 is provided as a separate file.

**Note S1: Clustering time series into characteristic trajectories**

We encoded the time series associated with each spacer as a string of characters, using letters A-F to indicate all possible combinations for the presence or absence of that particular spacer, its target, and its corresponding CRISPR lineage. For example, let us define A = [lineage, spacer, and target present]; B = [lineage and spacer present, target absent]; and C = [lineage present, spacer and target absent]. Then, a 3-sample time series in which the lineage was first observed alone, then the spacer and target appeared, and finally the target disappeared, will be codified by the string {CAB}. "Empty" states (that is, those characterized by the absence of spacer, CRISPR lineage, and target) were removed from both ends of the string and collapsed to a single instance if appearing more than once between non-empty states. For example, let us define F = [lineage, spacer, and target absent]; then, the string {FFCAFFFBF} will be reduced to {CAFB}. Such compression of empty states is justified by two reasons: First, the first and last sampling times do not correspond to any meaningful event, such as the onset of a treatment, and therefore there is no reason to assume that the dynamics should be synchronized among individuals. Because of that, empty states at the beginning or the end of the string are not informative. Second, because the interval between samples is relatively long (2 weeks), the number of consecutive empty states is not informative of the length of those absence periods. More precisely, lineages and targets could disappear and reappear several times without leaving any trace in the sampled time series. In consequence, empty states only inform us that the dynamics are intermittent. By collapsing internal empty states in the string, we preserve such information about intermittency in a more compact manner.

To cluster similar time series and obtain representative trajectories, we performed pairwise alignments of all strings using the following scores: 1 per match, -2 per mismatch, -2 per gap opening, and -1 per gap extension. These scores were chosen to prevent spurious pairing between short trajectories, that would be obtained if the penalties for mismatches and gap openings were not sufficiently large compared to the reward for matches. (Indeed, the combination [1,-1,-1,0] leads to a single cluster that contains all the time series, and [1,-1,-1,-1] produces three very large clusters encompassing 60% of all time series.) The gap extension penalty was included to avoid nesting short strings into longer strings. Note that, because internal empty states were collapsed during preprocessing, the length of a string is informative of the temporal prevalence of lineages and targets. Therefore, by penalizing gap extension, short trajectories that represent transitory infections can be distinguished from long trajectories that represent endemic or recurrent infections.

As shown in Figure S2, alternative scoring schemes produce similar clusters (in terms of the normalized variation of information) if they fulfil two conditions: (i) the penalty for mismatches and gap opening is greater than the reward for matches, and (ii) there is a cost for gap extension. If the cost of mismatches and gaps is too low, most time series become clustered in a few large (and poorly informative) clusters. If there is no cost for gap extension, the algorithm is unable to separate transitory from recurrent or endemic dynamics. The scoring scheme [2,-4,-4,-2] was included in Figure S2 to show that it produces almost identical results as [1,-2,-2,1]. The reason is that the network-based algorithm that we used for clustering (Infomap) is insensitive to the global scaling of weights. As a result, scoring schemes that differ by a constant multiplicative factor are fully equivalent in practice.

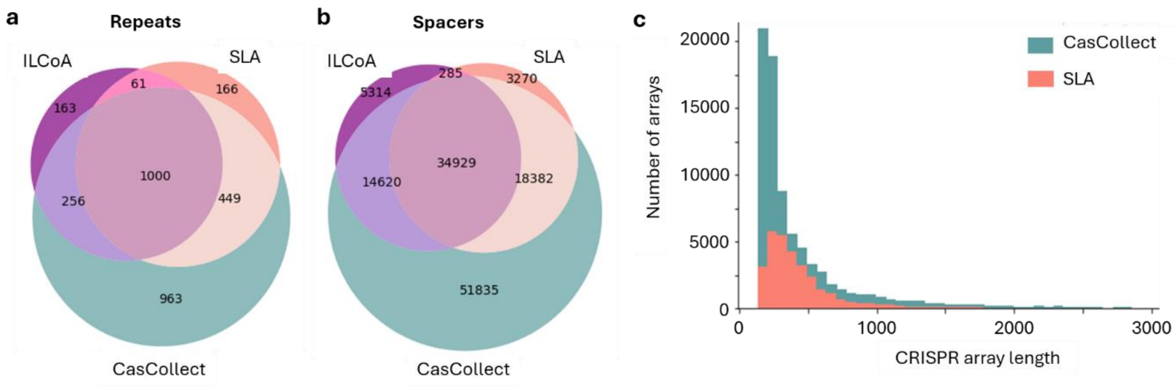


**Figure S1:** Comparison of CRISPR arrays obtained by CasCollect (followed by cleaning with CRISPRCasTyper) with those obtained by directly running CRISPRCasTyper on sample-level assemblies (SLA) and individual-level coassemblies (ILCoA). (a) Venn diagram showing the overlap in CRISPR repeats (clustered at 90% identity) among the three approaches. (b) Venn diagram showing the overlap in CRISPR spacers (clustered at 90% identity) among the three approaches. (c) Length frequency distribution of CRISPR arrays obtained with CasCollect and SLA.

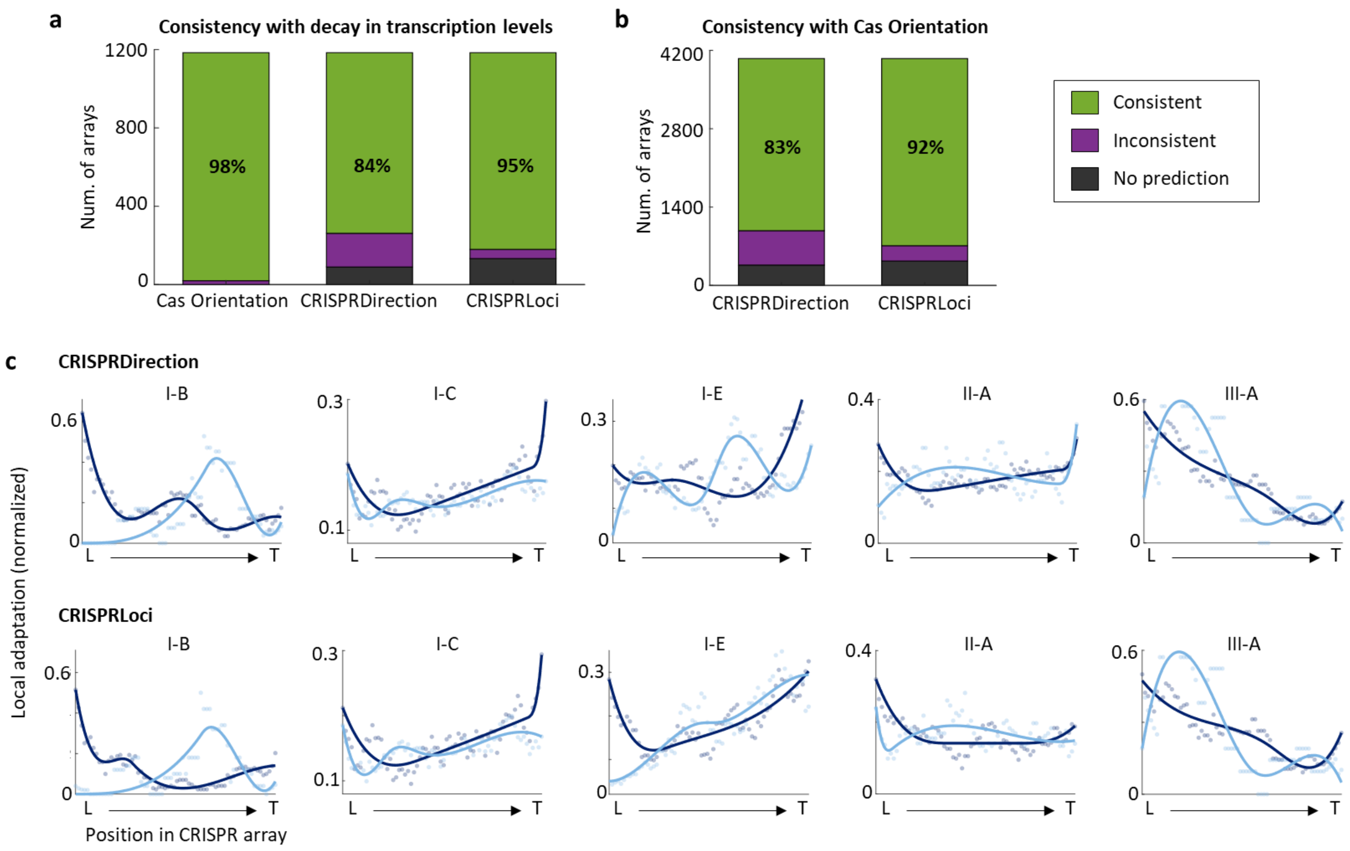


**Figure S2:** Robustness of orientation-based results with respect to the methods used to infer CRISPR orientation. (a) Consistency of predicted orientations with respect to the expected trend of reduced transcription in middle and distal regions of the array. (b) Consistency of CRISPRDirection and CRISPRLoci with respect to the predictions of CRISPR Orientation. CRISPRDirection and CRISPRLoci failed to predict the orientation of approximately 10% of the arrays in the proximity of Cas genes (black). The percentages in (a) and (b) indicate the fraction of consistent orientations with respect to all available predictions. (c) Relative position of locally adapted spacers within the CRISPR array. L: leading end; T: trailing end.

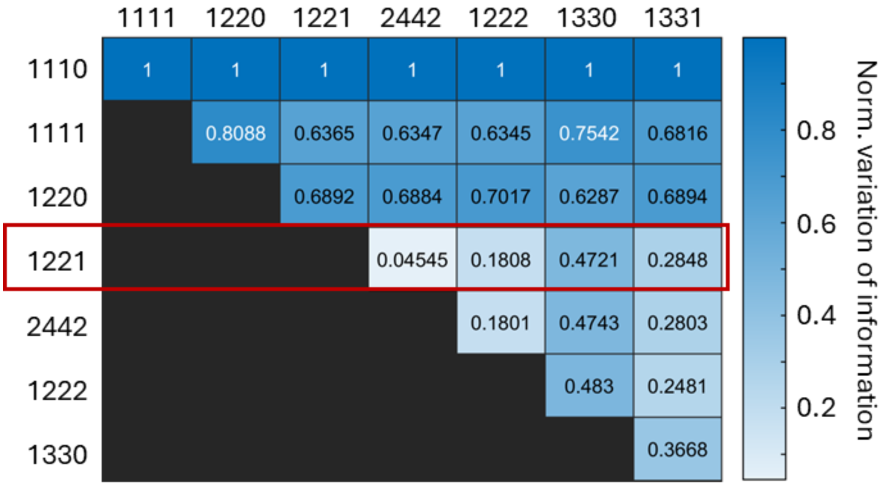


**Figure S3:** Comparison of the clustering of temporal dynamics obtained with different parameters. The normalized variation of information takes values from 0 (if two sets of clusters are identical) to 1 (if both sets are completely uncorrelated). Four-digit codes indicate the parameters used to align the temporal dynamics: The first digit corresponds to the reward per match, the second is the penalty for mismatch, the third is the penalty for gap opening, and the fourth is the penalty for gap extension. The red rectangle indicates the parameter combination used in the article.

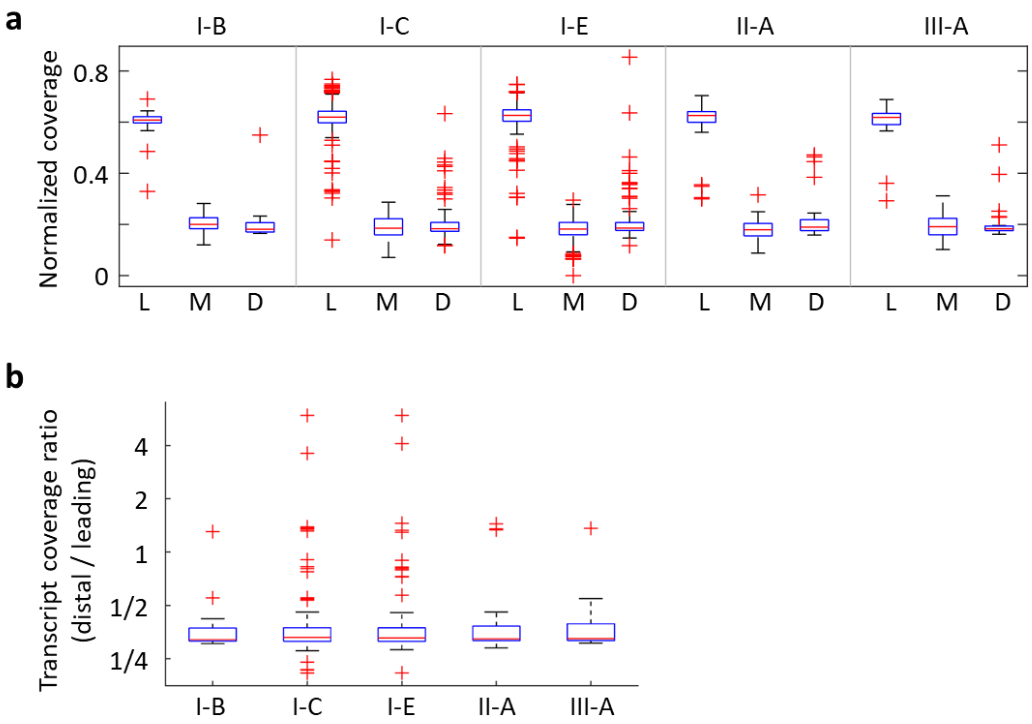


**Figure S4:** Differential expression of spacers as a function of their location within the CRISPR array. (a) Normalized transcript coverage of leading (L), middle (M), and distal (D) regions of the array. Each region was defined as one third of the total array length. To compare among arrays with different overall expression levels, transcript coverages were separately normalized in each array, so that the total coverage per array was equal to 1. (b) Transcript coverage ratio, indicating the relative expression of the distal third of the array with respect to the leading third.

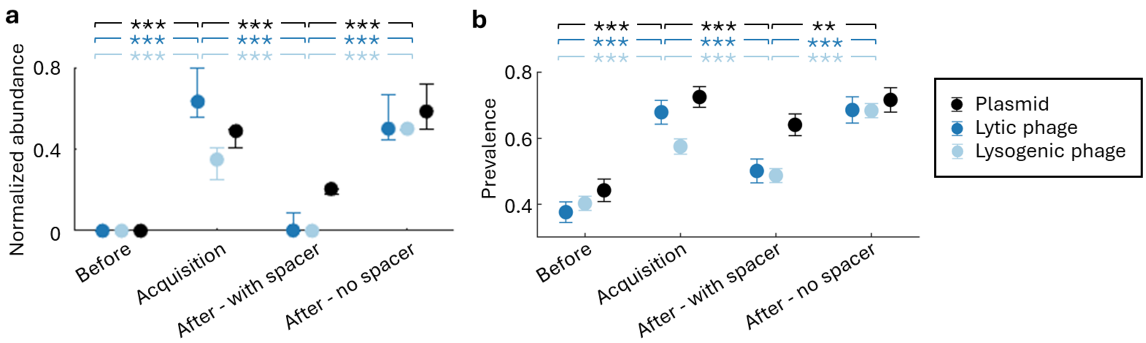


**Figure S5:** Effect of CRISPR-Cas immunity on MGE abundance and host diversity (see also Fig. 4). To control for possible biases due to different lineage sizes, for each condition (see below), we grouped spacers by lineage and calculated their median normalized abundance and mean prevalence. As a result, each lineage contributed with at most one data point to each condition. (a) Median normalized abundance of targeted MGE (plasmids, lytic/non-lysogenic phages, and lysogenic phages) before the spacer was first observed in the host population ("Before"), in the sample in which the spacer was observed for the first time ("Acquisition"), in subsequent samples in which the spacer was present ("After – With spacer"), and in subsequent samples that contained the same CRISPR lineage but not the spacer ("After – No spacer"). Error bars indicate the 95% confidence intervals for the median, based on 2,000 bootstraps. All comparisons shown above the plot are statistically significant with $p$ < 10-4 (Mann-Whitney test). (b) Mean prevalence of targeted MGE, calculated as the fraction of samples in which the target is detected. Error bars indicate the 95% confidence intervals for the mean. Statistical significance based on Student's T test: $p < 0.05$ (*), $p < 0.01$ (**), $p < 0.001$ (***).

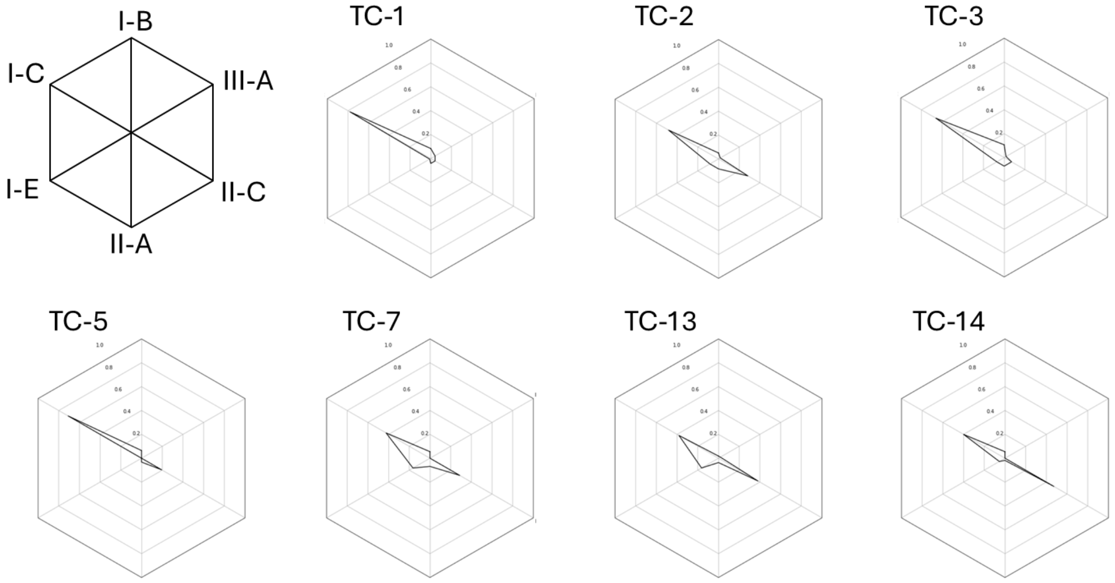


**Figure S6:** Relative representation of different CRISPR-Cas subtypes in the representative trajectory clusters (TC) shown in Figure 5.

**Table S1.** Comparison of CRISPR repeat and spacer diversity retrieved through three different methods. SLA: Sample-level assembly (from HMP2-IBD, MEGAHIT assembler) followed by CRISPRCasTyper. ILCoA: Individual-level coassembly (SPAdes) followed by CRISPRCasTyper. CasCollect: CasCollect (sample-level targeted assembly of CRISPR-Cas regions) followed by CRISPRCasTyper. See Methods for details.

| | SLA | | ILCoA | | CasCollect | |
|---|---|---|---|---|---|---|
| | Repeats | Spacers | Repeats | Spacers | Repeats | Spacers |
| Total | 30751 | 200073 | 8194 | 80358 | 79475 | 534232 |
| Clustered 90% | 1431 | 56390 | 1221 | 54960 | 2555 | 119411 |
